# Heartfelt Face Perception via the Interoceptive Pathway – an MEG study

**DOI:** 10.1101/200980

**Authors:** Jaejoong Kim, Hyeong-Dong Park, Ko Woon Kim, Dong Woo Shin, Sanghyun Lim, Hyukchan Kwon, Min-Young Kim, Kiwoong Kim, Bumseok Jeong

## Abstract

The somatic marker hypothesis proposes that the cortical representation of visceral signals is a crucial component of emotion processing. No previous study has investigated the causal relationships among brain regions that process visceral information during emotional perception. In this magnetoencephalography study of 32 healthy subjects, heartbeat-evoked potentials (HEPs), which reflect the cortical processing of heartbeats, were modulated by the perception of a sad face. The modulation effect was localized to the prefrontal cortices, the globus pallidus, and an interoceptive network including the right anterior insula (RAI) and anterior cingulate cortex (RACC). Importantly, our Granger causality analysis provides the first evidence for increased causal flow of heartbeat information from the RAI to the RACC during sad face perception. Moreover, this HEP modulation effect was neither an artefact nor an effect of visual-evoked potentials. These findings provide important progress in the understanding of brain-body interactions during emotion processing.

## Introduction

According to the James-Lange theory and the somatic marker hypothesis, emotional feelings are the mental experience of bodily states (A. Damasio & Carvalho, 2013; James, 1884). More specifically, emotional stimuli usually induce a change in bodily status (Williams et al., 2005). Then, various feelings subsequently emerge from the perception of bodily status, including a sense of viscera (A. Damasio & Carvalho, 2013). Many previous studies have shown evidence of emotional stimulus-evoked somatic responses. For example, a fearful stimulus modulates autonomic responses such as heart rate and skin conductance (Critchley et al., 2005; Williams et al., 2005), and a disgusting stimulus induces tachygastria (Harrison, Gray, Gianaros, & Critchley, 2010). Moreover, neuroimaging studies using functional magnetic resonance imaging (fMRI) or electroencephalograms (EEGs) have shown that the generation of bodily responses by an emotional stimulus is related to the activation of subcortical regions such as the amygdala and hypothalamus (A. R. Damasio et al., 2000). On the other hand, there is little evidence that supports a change in cortical interoceptive processing in the brain when experiencing emotions. Given that the signals from internal organs cannot be identified without explicit measuring devices such as electrocardiograms (ECG), it is difficult to investigate the brain activity that is directly evoked by interoceptive signals. Thus, previous studies have explored changes in interoceptive processing during the experiencing of emotional states and reported relationships with the anterior insular and anterior cingulate cortical activity (Adolfi et al., 2016; Critchley et al., 2005). Heartbeat-evoked potentials (HEPs), which are obtained by averaging electrophysiological signals time-locked to heartbeats, have been reported to be associated with heartbeat perception accuracy (Pollatos & Schandry, 2004), suppressed by pain perceptions (Shao, Shen, Wilder-Smith, & Li, 2011), and modulated by empathy feelings (Fukushima, Terasawa, & Umeda, 2011). HEP amplitudes are also attenuated in mood-related psychiatric disorders, including depression (Terhaar, Viola, Bär, & Debener, 2012) and borderline personality disorder (Müller et al., 2015), suggesting a potential link between HEPs and aberrant emotional processing. Moreover, several recent studies have successfully shown using HEPs that cortical interoceptive processing is modulated by emotional processing. One study reported that HEPs are modulated by emotional arousal (Luft & Bhattacharya, 2015). An another study reported that HEPs in infants are modulated by fearful and angry emotional facial expression video clips (Maister, Tang, & Tsakiris, 2017). Finally, one study, using high density EEGs and natural affective scenes, localized the source of HEPs to a frontal-insular-temporal network, including the anterior insula and anterior cingulate cortex (Couto et al., 2015). However, the modulation of directional interoceptive information flow between these regions during emotion processing, which is predicted by the somatic marker hypothesis (A. Damasio & Carvalho, 2013), has not been investigated yet. Therefore, in this study, we aimed to identify the modulation of directional interoceptive information flow during emotion perception using HEPs. To test this idea, we used emotional faces and emotional emoticons that conveyed text-based emotions to evoke an emotional feeling while measuring HEPs with magnetoencephalography (MEG). To verify the precise source of HEP modulation, T1-weighted structural MRI was performed for all subjects. Importantly, we applied Granger causality (GC) analysis (Barnett & Seth, 2014) to sources to identify information flow between the sources of HEP modulations.

We formulated the following specific hypotheses. First, we expected that HEPs would be modulated by emotional expression and that this effect would be seen by different spatiotemporal dynamics between emotional and neutral stimulus presentations. Specifically, we used sad and happy expressions as emotional stimuli to observe a HEP modulation effect after visual stimulus presentation because these emotional stimuli, as well as neutral faces, are known to be less influenced by the timing of stimuli presentations within cardiac cycle (Garfinkel & Critchley, 2016). Although emotional expression is not verified by this method, we expected that emoticons with these emotions (sad and happy) and neutral emoticons would also be less influenced by the timing of stimuli presentations within cardiac cycle.

Second, we expected that the modulation of HEPs by emotional expressions would be localized in previously known convergence regions of interoception and emotion, such as the anterior insula and anterior cingulate cortex, when using source level analysis (Adolfi et al., 2016). Third, we expected that information flow between these interoceptive regions would be modulated by emotional expression. To be more specific, we expected that bottom-up heartbeat information processing starting from the anterior insula, which re-represents the viscerosensory information from the posterior insula, to the anterior cingulate cortex would be enhanced by emotional expressions (Medford & Critchley, 2010). This pathway is hypothesized to be involved in the generation of emotional feelings and the processing of the subjective salience of emotions using interoceptive signals (Medford & Critchley, 2010; Smith & Lane, 2015; Uddin, 2015). Finally, we hypothesized that neural activity evoked by heartbeats would show spatiotemporal patterns different from the cortical activity evoked by visual stimuli. That is, we expected that the MEG signals time-locked to the cardiac cycle onset would cause the visually evoked effect to disappear and vice versa.

## Methods

### Participants

Forty healthy participants (19 females, mean age of 24.03 ± 3.28 years) volunteered for this experiment. The expected effect sizes were not known in advance, so we chose a sample size of approximately 40 participants, which was approximately two times more than that of previous MEG and EEG studies of HEPs (Babo-Rebelo, Richter, & Tallon-Baudry, 2016; Fukushima et al., 2011; H.-D. Park, Correia, Ducorps, & Tallon-Baudry, 2014).

MEG recordings consisting of 4 runs were completed in one visit. High-resolution T1 weighted MRI scans were acquired at another visit. In this MRI session, all subjects underwent both functional MRI experiments, consisting of emotional discrimination (unpublished data) and/or decision tasks (unpublished data), and other structural MRI scans, such as diffusion tensor imaging (unpublished data). We failed to acquire MEG data for five of the forty subjects due to magnetic field instability. Another three subjects were excluded during analysis because their ECG data were too noisy or absent. Therefore, thirty-two subjects were included for further analysis.

A structured interview was conducted using the Korean version of the Diagnostic Interview for Genetic Studies (Joo et al., 2004). None of the subjects had current neurological or psychological diseases. Subjects completed the Patient Health Questionnaire-9 (PHQ-9)(Kroenke, Spitzer, & Williams, 2001), a self-rating scale for depression, and the Korean version of the Verbal Abuse Questionnaire (K-VAQ) (Jeong et al., 2015) because we suspected that previous emotional abuse history might affect responses to emotional stimuli. The mean PHQ-9 score was 3.62 (SD: 2.59, n = 39). The mean score of the peer verbal abuse (PeVA) and parent verbal abuse (PaVA) assessments were 22.61 (SD: 9.65, n = 38) and 21.95 (SD: 7.82, n = 38), respectively. All thirty-two participants included in the analysis completed the PHQ-9 (mean and SD: 3.68 and 2.69, n = 32), while two participants did not complete the K-VAQ (mean and SD of PeVA: 21.73 and 7.26, mean and SD of PaVA: 23.43 and 10.29, n = 30). All participants provided written informed consent to participate in the experiment. The study was approved by the Korean Advanced Institute of Science and Technology Institutional Review Boards in accordance with the Declaration of Helsinki.

### Standardization of emotional stimuli

Stimuli consisted of forty-five emotional faces and forty-five text-based emotional emoticons. Forty-five faces expressing happy, sad, and neutral emotions were selected from the Korean Facial Expressions of Emotions (KOFEE) database (J. Y. Park et al., 2011). Text-based happy and sad emoticons were searched for on the world-wide web. Then, we created scrambled emoticons that did not have configurable information and used these scrambled emoticons as neutral condition emoticons; examples of text-based emoticons are provided in the supplementary materials (Fig. S1). Ninety emotional expressions, including faces and text-based emoticons, were standardized in independent samples consisting of forty-seven healthy volunteers (21 females, mean age of 28.43 ± 4.31 years). These participants were asked to rate the intensity of the feeling they felt towards the emotional expressions of 90 stimuli (45 faces and 45 facial emoticons with happy, neutral, and sad emotions) on an 11-point Likert scale (−5 to +5). We compared the mean absolute value of the four emotional expressions and two neutral expressions, which we called ‘feeling intensity’ or ‘emotionality’ (Citron, Gray, Critchley, Weekes, & Ferstl, 2014). A repeated-measures analysis of variance (repeated-measures ANOVA) with a 2 stimuli (face, emoticon) by 3 valences (happy, sad, neutral) design was performed on the means and variances of emotionality score independently. In the repeated-measures ANOVA of the means, there was a significant main effect of valence (F (1.744, 80.228) = 272.618, p < 0.001, Greenhouse-Geisser-corrected), while there were no differences between emoticons and faces (F (1, 46) = 0.011, p = 0.919) and no interaction between those two main effects (F (1.685, 77.488) = 0.285, p = 0.818, Greenhouse-Geisser-corrected). In addition, a post hoc *t* test revealed that there was no difference between sad and happy conditions (p = 0.082) but a significant difference between emotional and neutral conditions (p < 0.001 for both sad and happy compared with neutral). In the repeated-measures ANOVA of the variance, the variance for the emoticons was significantly larger than the variance for the faces (F (1, 46) = 16.108, p < 0.001), while there was no difference in the variance between emotions (F (1.268, 58.342) = 2.608, p = 0.079, Greenhouse-Geisser-corrected) and no significant interaction (F (1.347, 61.963) = 4.831, p = 0.066, Greenhouse-Geisser-corrected). Additionally, participants from the main experiments also performed the rating procedure above before MEG recordings.

### MEG experimental task

During the MEG recording, ninety stimuli consisting of 45 faces and 45 text-based emoticons were presented in the centre of a screen using in-house software, the KRISSMEG Stimulator 4. The size, duration, and stimulus onset asynchrony (SOA) of all the stimuli were 27×18 cm, 500 ms and 1500 ms, respectively, and the order of stimuli presentation was pseudo-randomized. Participants completed 4 runs, and each run contained 180 stimuli (30 sad faces, 30 happy faces, 30 neutral faces, 30 sad emoticons, 30 happy emoticons, 30 neutral emoticons each) and lasted 270 s. In addition, to maintain participants’ attention to task, the participants were instructed to discriminate between sad and happy by pressing a button when a question mark appeared. The question mark randomly appeared on the screen every 9 to 15 trials (mean = 11.3). A total of 4.7% of the trials were response trials. The results of the discrimination task were not used in further analyses.

### Acquisition

A 152-channel MEG system (KRISS MEG, Daejeon, Korea, 152 axial first-order double-relaxation oscillation superconducting quantum interference device (DROS) gradiometers) covering the whole head was used for MEG recordings in a magnetically shielded room for 60–90 minutes at a sampling rate of 1,024 Hz. The relative positions of the head and the MEG sensors were determined by attaching four small positioning coils to the head. The positions of the coils were recorded at intervals of 10–15 min by the MEG sensors to allow co-registration with individual anatomical MRI data. The maximum difference deviations between head positions before and after the run were < 2 mm, and the goodness of fit (GoF) > 95%. EEGs of eye and muscle artefacts were recorded simultaneously with the MEG recordings. During MEG recordings, participants were seated with their heads leaned back in the MEG helmet. The translation between the MEG coordinate systems and each participant’s structural MRI was made using four head position coils placed on the scalp and fiducial landmarks (Hämäläinen, Hari, Ilmoniemi, Knuutila, & Lounasmaa, 1993).

### Data preprocessing

Data were processed with the FieldTrip toolbox (Oostenveld, Fries, Maris, & Schoffelen, 2011). First, raw data were epoched from 700 ms before stimulus onset to 1300 ms after stimulus onset. Epochs containing large artefacts were rejected by visual inspection. After artefact trial rejection, eye movement and cardiac field artefacts (CFA) were removed by independent component analysis (ICA) (Hyvärinen, Karhunen, & Oja, 2004) using the function “ft_componentanalysis” with the runICA algorithm. Full epochs were submitted for ICA, and twenty components were identified for each of the six conditions. Two neurologists visually inspected each component, and the components that showed typical spatial and temporal patterns of CFA, eye blinking and movement noise were removed. On average, 2.44 ICs were removed per subject, including one or at most two ICs reflecting CFA (in most cases, one IC reflecting CFA was detected) After removing noise, the data were filtered with a 1-40 Hz Butterworth filter. Then, the HEPs for each stimulus condition were extracted by subsequent epoching, which was time-locked to the R peak of every epoch. R peaks were detected using the Pan-Tompkins algorithm (Pan & Tompkins, 1985), and the HEPs of each condition were extracted by epoching 500 ms before the R peak to 600 ms after the R peak in the epoch of each condition. Because a heartbeat enters the central nervous system (CNS) approximately 200 ms after the R peak by vagal afferent stimulation at the carotid body (Eckberg & Sleight, 1992) and a visual stimulus enters the CNS immediately through the retina, a heartbeat that occurs before a −200 ms visual stimulus onset stimulates the brain earlier than a visual stimulus onset. Therefore, we excluded R peaks that occurred before 200 ms of a stimulus onset to include heartbeat-evoked processing that occurred only after a visual stimulus. Therefore, this procedure excluded the cortical input of a heartbeat that occurred before the visual stimulus. The R peaks after 700 ms of stimulus onset were also excluded because that HEP epoch would contain the next visual stimulus onset. The mean number of HEP epochs per conditions after the HEP extraction procedure was 102.78, and there was no significant difference in number of epochs between conditions (one-way repeated-measures ANOVA, F (3.22, 100.04) = 0.29, p = 0.84, Greenhouse-Geisser-corrected). Finally, a baseline correction was performed using a pre-R-peak interval of 300 ms, and trials of the same condition for each subject were averaged. We expected that HEP modulation induced by brief emotional faces and emoticons used in this study would be much smaller than the CFA, especially considering that sad and happy faces are typically considered to induce low arousal (Liu, Chen, Hsieh, & Chen, 2015). It is known that HEP amplitudes and electrocardiac fields overlap up to 450 ms after the R peak (Dirlich, Vogl, Plaschke, & Strian, 1997). Therefore, to avoid our effect of interest being obscured by CFA, we attempted to maximally control the CFA by using the time window of 455-595 ms after each R peak for the statistical analyses of HEPs; this time window is known to have a minimal influence on CFA and, thus, has been used in many previous studies (Gray et al., 2007; Müller et al., 2015; Schulz et al., 2015).

### Sensor analysis: Cluster-based permutation paired *t* tests between each emotional condition and the neutral condition

We compared the HEPs of an emotional condition and a neutral condition. Four tests were performed, including a sad face vs. a neutral face, a happy face vs. a neutral face, a sad emoticon vs. a neutral emoticon, and a happy emoticon vs. a neutral emoticon. To address multiple comparison problems, we used cluster-based permutation paired *t* tests. These tests were performed as follows. First, data were downsampled to 256 Hz to make the computation efficient, and paired *t* tests were performed at all time points between 455 and 595 ms and all sensors. Then, significant spatiotemporal points of uncorrected p-values below 0.05 (two-tailed) were clustered by spatiotemporal distance and the summed t-value of each cluster was calculated. After calculating the cluster t-stat, a permutation distribution was created by switching condition labels within subjects randomly, calculating the t-value of the paired *t* test between permutated conditions, forming clusters as mentioned above, selecting the maximum cluster t-value and repeating this procedure 5000 times. Finally, after the maximum cluster t-values of each permutation created a permutation distribution, the corrected p-value original clusters were calculated. Additionally, because many previous studies of HEPs have also reported a HEP effect in earlier time windows (Couto et al., 2015; Fukushima et al., 2011; Luft & Bhattacharya, 2015; Pollatos & Schandry, 2004), we performed additional cluster-based permutation paired *t* tests in earlier time windows between conditions that showed significant differences in the original time window of 455-595 ms post-R peak as a control analysis. The time windows of 315-455 ms post-R peak and 175-315 ms post-R peak (which have the same length of the original time window) were tested separately. Note that the results of this control analysis might have been influenced by the residual CFA (which might be smaller in the 315-455-ms window than in the 175-315-ms window).

### Source analysis

Source reconstruction was conducted using the MATLAB package Brainstorm (Tadel, Baillet, Mosher, Pantazis, & Leahy, 2011). To estimate the time courses of both cortical and subcortical activity, we used the default settings in the open source MATLAB toolbox Brainstorm’s implementation of the deep brain activity model using the minimum norm estimate (MNE) (Attal & Schwartz, 2013; Tadel et al., 2011). First, cortical surfaces and subcortical structures, including the amygdala and basal ganglia, were generated for each subject from 3T MPRAGE T1 images using FreeSurfer (Fischl, 2012). The individual heads/parcellations were then read into Brainstorm (Tadel et al., 2011) along with tracked head points to refine the MRI registration. In Brainstorm, a mixed surface/volume model was generated, and 15,000 dipoles were generated on the cortical surface with another 15,000 dipoles generated in the subcortical structure volume. Refining the registration with the head points improves the initial MRI/MEG registration by fitting the head points digitized at the MEG acquisition and the scalp surface. Using the individual T1 images and transformation matrix generated as above, a forward model was computed for each subject using a realistic overlapping spheres model. The source activity for each subject was computed using the MNE (Brainstorm default parameters were used). The source map was averaged over the time window of 488 ms to 515 ms, which showed a significant difference between a sad face and a neutral face at the sensor level (other emotional conditions were not significantly different from neutral conditions in cluster-based permutation paired *t* tests). Then, this averaged spatial map was exported to SPM12 software (Penny, Friston, Ashburner, Kiebel, & Nichols, 2011), and subsequent statistical tests were performed. Paired *t* tests were used to identify regions that had a different HEP time course within the selected time window between sad and neutral faces. In source analysis, data were downsampled to 512 Hz using a sufficient number of time points in the GC analysis.

### GC analysis of HEP source activity

After identifying brain regions that had different time courses, we performed a GC analysis (Barnett & Seth, 2014) on two regions of interest, the right anterior insula (RAI) and the right anterior cingulate cortex (RACC), to determine whether the effective connectivity between these regions is modulated differently in emotional conditions compared to a neutral condition. These regions are known to be core regions of interoceptive processing and feelings according to many previous studies (Adolfi et al., 2016; A. R. Damasio, 2003; Smith & Lane, 2015). Moreover, emotion processing in the AI and ACC is known to be right lateralized, especially in the AI (Craig, 2009; Gu, Hof, Friston, & Fan, 2013). The time courses of these two regions were extracted from voxels that showed a significant difference in source analysis. Detailed information, including the coordinates of the ROIs, are provided in the supplementary materials.

In GC (Granger, 1988) analysis, time course Y causes time course X if the past time points of Y and X explain X better than the past time points of X alone. The GC is formulated by the log-likelihood ratio between the residual covariance matrix of the model that explains X by the past of X and Y and the residual covariance matrix of the model that explains X by the past of X alone (Barnett & Seth, 2014).

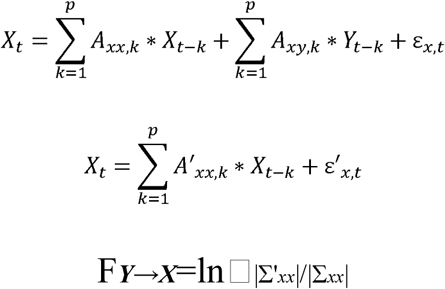

A is the matrix of the regression coefficient, epsilon is the residual, sigma is the covariance matrix of the residual, and F is the GC of X and Y. All calculations were performed using a multivariate GC toolbox (MVGC toolbox) (Barnett & Seth, 2014).

The time courses of two ROIs were extracted for every trial for each subject. To satisfy the stationarity assumptions of GC analysis, we used short time window GC estimations with sliding windows (Ding, Bressler, Yang, & Liang, 2000; Seth, Barrett, & Barnett, 2015). This approach was also appropriate for our analysis of the HEP modulation effect, which was identified in a very short time window and thus required a high temporal precision of the GC estimation. The size of the window was 60 ms, the step size was 2 ms (Cohen, 2014; Ding et al., 2000), and the GC calculation was performed for the whole epoch, which started from the [−500 ms −440 ms] window to the [540 ms 600 ms] window. Stationarity was further controlled by removing the average event-related potential from each trial (Wang, Chen, & Ding, 2008). The model order was determined using Akaike information criterion (AIC) to a maximum order of seven time points, which corresponded to 14 ms. After model estimation, we tested the stationarity of the model by examining whether the spectral radius ρ(A) > 1 in every time window and every subject (Barnett & Seth, 2014). Although we tried to control every time window to satisfy the stationarity assumption, after the [507 ms 567 ms] time window, 21 participants violated the stationarity assumption. Additionally, violations occurred at approximately 0 ms post-R peak in three participants and at 370 ms (around the T peak) in one participant. There was a similar pattern even when we tested variable lengths of the time window and model order. We suspected that this stationarity violation might be induced by CFAs. Therefore, we again used the time window starting from the [455 ms 515 ms] window to the [507 ms 567 ms] window, and every time window satisfied the stationarity assumption in every participant. Pairwise GC analysis of the two ROIs (two GCs - RAI to RACC, RACC to RAI) were performed for emotional and neutral conditions. To compare between emotional and neutral conditions, GC baseline normalization was performed in both conditions by calculating the change in the GC relative to the average GC between the [−330 ms −270 ms] window and the [−130 ms −70 ms] window (Cohen, 2014). Time windows approximately 0 ms post-R peak were not used as a baseline because three subjects violated the stationarity assumption. Finally, the 2 estimated GCs of emotional and neutral conditions were compared using a cluster-based permutation paired *t* test for all time windows starting from the [455 ms 515 ms] window to the [507 ms 567 ms] window (Oostenveld et al., 2011). Therefore, the multiple comparisons were controlled for the number of GCs and the number of time windows using a cluster-based permutation paired *t* test.

### Analysis of physiological data

#### Heartbeat distributions in each condition

To show that the HEP effect was not the result of a biased heartbeat distribution, within the original visual epoch, we divided the visual epoch between −200 ms and 700 ms, i.e., the beginning and end of the HEP epoching, into 100-ms time windows (a total of nine time bins) and counted how many heartbeats there were in these time windows. We completed this procedure for every condition. Then, ANOVA between the nine time bins was performed to test whether the occurrence of heartbeats was the same in every time bin.

#### Mean interbeat interval (IBI) and IBI modulation in each condition

IBI was calculated by measuring the time interval between two consecutive R peaks. The average IBI of all subjects was 921 ± 100 ms, with a range of 789 ms to 1152 ms. Moreover, we calculated the IBI of all conditions by measuring the IBI between the R peak immediately after the stimulus onset and the R peak immediately following that one and averaging these IBIs separately for each condition. Then, a one-way repeated-measures ANOVA of IBIs between every condition was performed to test whether the IBIs were different across conditions.

### Analysis to exclude the effects of visual processing

#### Surrogate R peak analysis

To test whether the HEP modulation effect was time-locked to the heartbeat, we created 100 surrogate R peaks that were independent of original heartbeats (H.-D. Park et al., 2016; H.-D. Park et al., 2014). Surrogate R peaks were created by randomly shifting the original R peaks (−500 ms ∼ +500 ms) the same amount in each subject (H.-D. Park et al., 2016). Then, we computed the surrogate HEPs with surrogate R peaks and performed the same cluster-based permutation *t* tests between the conditions that showed a significant difference in the sensor level analyses. Finally, we calculated the distribution of the maximum cluster statistics of the surrogate R peaks and the calculated position of our original cluster statistics in this distribution to show that the heartbeat-locked effect was significantly larger in such a distribution.

#### Analysis of visual-evoked potential (VEP)

To test whether the HEP modulation effect was confounded by a VEP effect, we performed the same cluster-based permutation test with a visual stimulus-locked potential. The time window of 0 ms to 1300 ms after stimulus onset was used as test input. We compared the topology of the significant clusters between HEPs and VEPs at the sensor level. Then, we performed source localizations of VEP activity in significant cluster time windows (78-198 ms and 815-948 ms separately) and exported them to SPM12 to perform statistical tests between emotional and neutral conditions with the same methods used in the HEP analysis (including the absolute value differences). Then, we compared the resulting sources with the results of the HEP analysis. This VEP analysis was not performed for all conditions but instead for those conditions that had a significant HEP modulation effect so that we could compare the significant modulation of HEPs with the VEP effect.

## Results

### Sensor analysis

#### Significant difference in HEPs between sad face and neutral face perceptions

A HEP cluster showing a significant difference between the perception of a sad face and of a neutral face was found in the right frontocentral sensors within a 488 ms-515 ms time range (Monte-Carlo p = 0.046, Fig. 2). In other conditions, including happy face vs. neutral face (Monte-Carlo p = 0.396), sad emoticon vs. neutral emoticon (Monte-Carlo p = 0.857), and happy emoticon vs. neutral emoticon (Monte-Carlo p = 0.710), there were no clusters showing significantly different HEP amplitudes. In the control analysis of the earlier time windows for the sad face condition (which showed significant difference), no significant clusters were found in either the 175-315 ms (Monte-Carlo p = 0.508) or 315-455 ms (Monte-Carlo p = 0.151) time windows. Note that we performed additional analysis on the sad emoticon condition, which is also a sad emotional expression similar to the sad face, but no HEP modulation effect was found. Detailed information is provided in the supplementary materials.

**Fig. 1.**
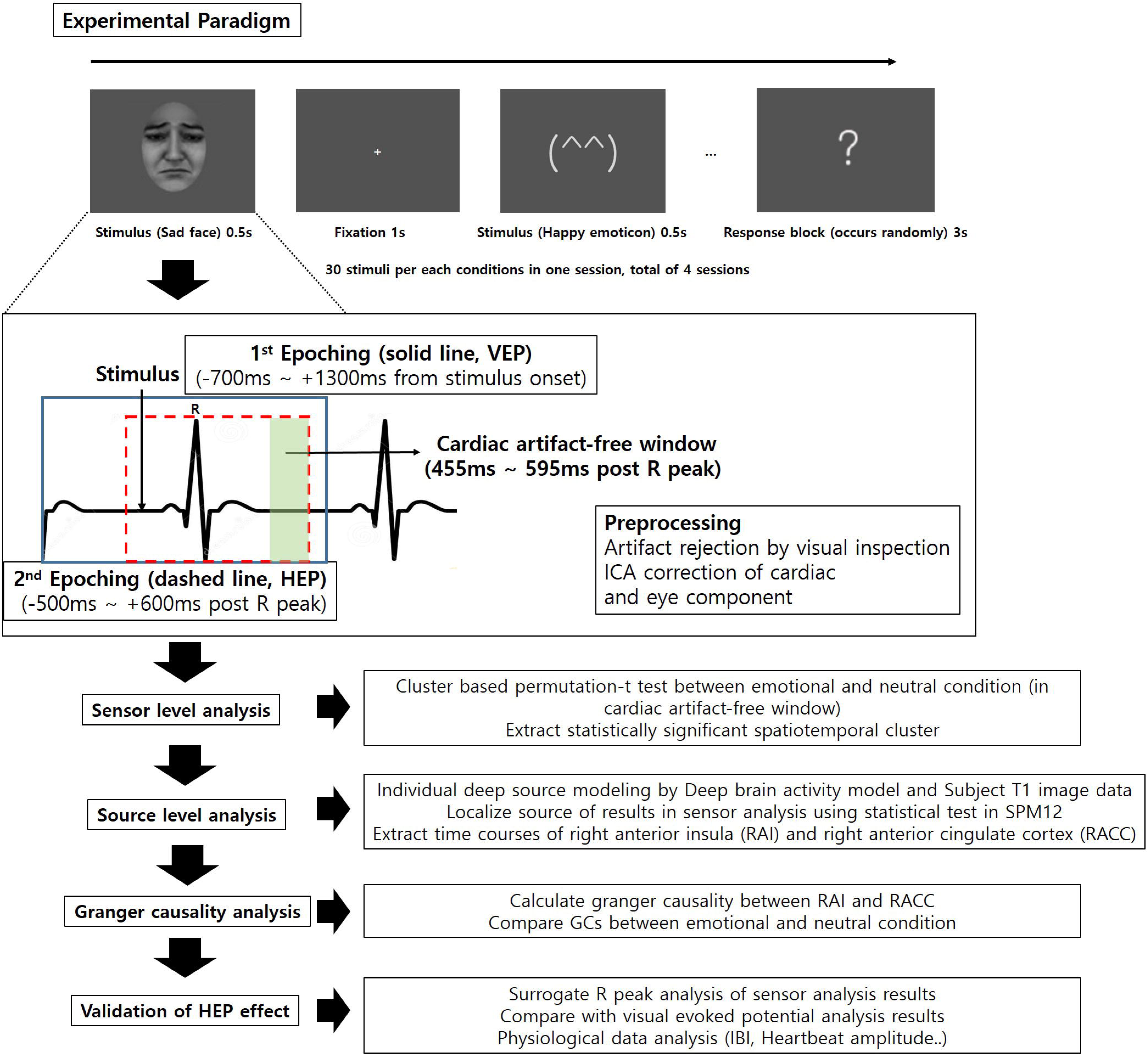
Overall experimental flow. The sad face used in this figure is different from the one used in the experiment (the sad face in this figure was generated by the FaceGen Modeller (http://www.facegen.com/)).

**Fig. 2.**
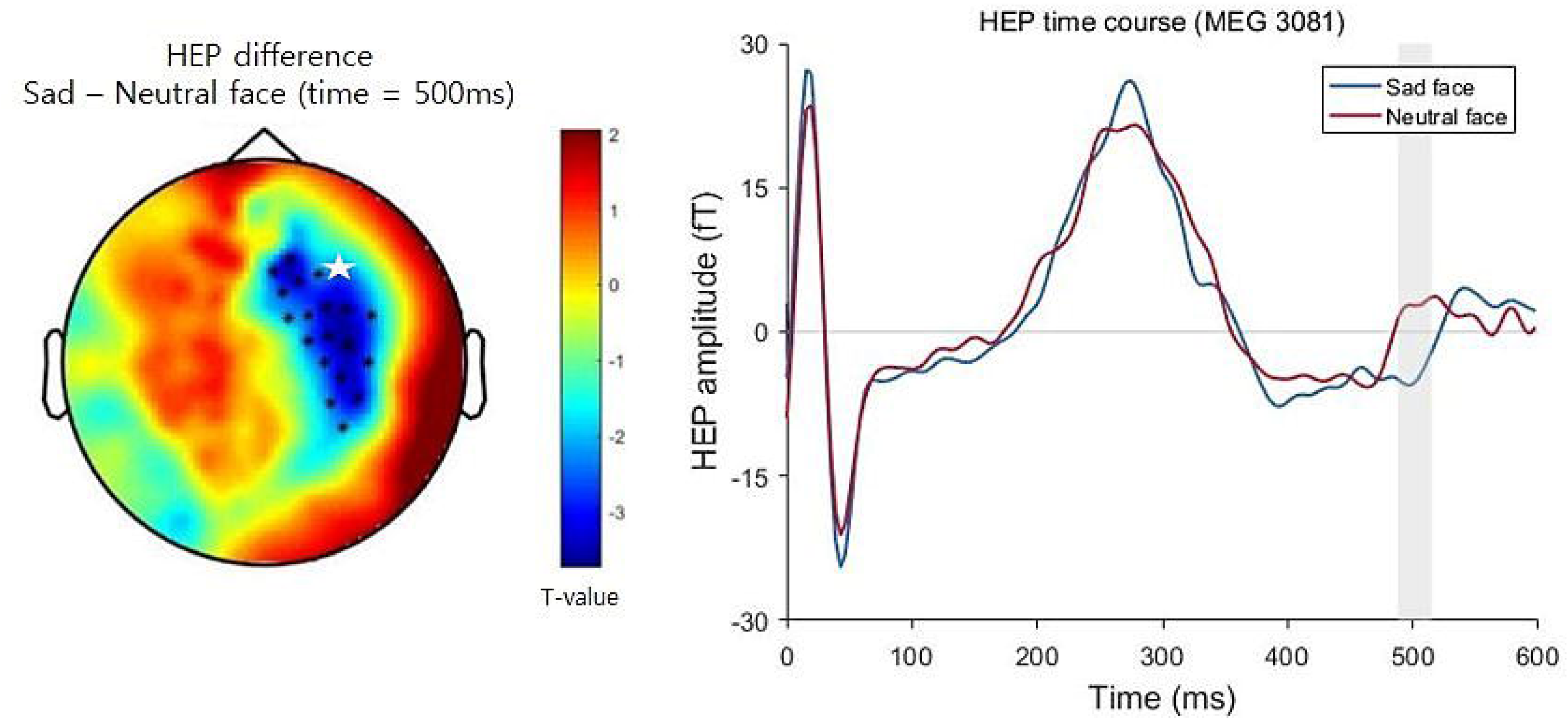
Topographic map (left) of the differences in HEPs between sad and neutral face conditions. Clusters showing significant differences are marked with a black dot (Monte-Carlo p < 0.05, cluster-corrected). In a single channel plot of a significant cluster (right), the shaded area represents the cluster time window that shows a significantly different time course between conditions. The channel plotted in the right figure is marked with a white star in the left topographic map.

### Source analysis

#### Interoceptive and prefrontal-basal ganglia networks as sources of HEP modulations to sad faces

With the cluster-forming criteria of p-value = 0.01 and 10 adjacent voxels, several regions that had different HEP time courses between sad face and neutral face conditions were identified. Four significant clusters appeared (Table 1). Briefly, these clusters included the right prefrontal cortices, the anterior insula, the anterior cingulate cortex, and the basal ganglia. More specifically, the first cluster (red in Fig. 3, p = 0.003, cluster-level family-wise error (FWE)-corrected) included the right prefrontal regions that consisted of the right superior frontal gyrus (RSFG), which is close to the dorsomedial prefrontal cortex (dmPFC), and the middle frontal gyrus (RMFG), which corresponded to the dorsolateral prefrontal cortex (dlPFC). The second cluster (green in Fig. 3, p = 0.001, cluster-level FWE-corrected) included the RAI and the right putamen (RP). The third cluster (blue in Fig. 3, p = 0.003, cluster-level FWE-corrected) included the RACC and the left anterior cingulate cortex (LACC). Finally, the fourth cluster (pink in Fig. 3, p < 0.001, cluster-level FWE-corrected) included the right basal ganglia, which consisted of the right globus pallidus (RGP) and the RP. Previous studies have reported the ACC and AI as sources of HEPs (Couto et al., 2015; H.-D. Park et al., 2017; H.-D. Park et al., 2014).

**Fig. 3.**
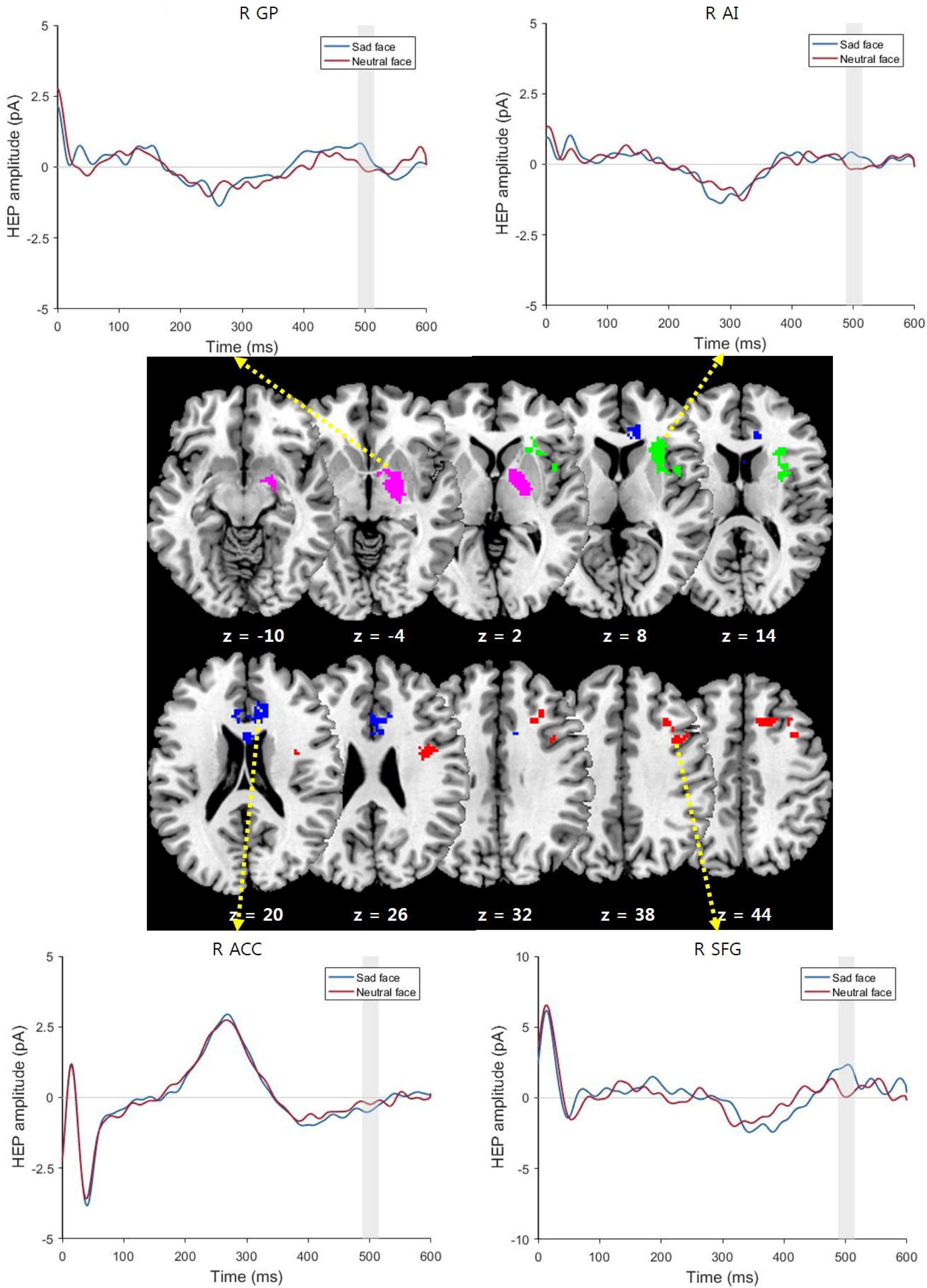
Brain regions showing significant differences in HEP modulations in the contrast of sad vs. neutral face conditions (p < 0.01, cluster-level FWE-corrected) and the mean time courses of HEPs from the 25 voxels surrounding the peak voxels. The yellow dashed arrow connects the cluster region and its corresponding time course.

**Fig. 4.**
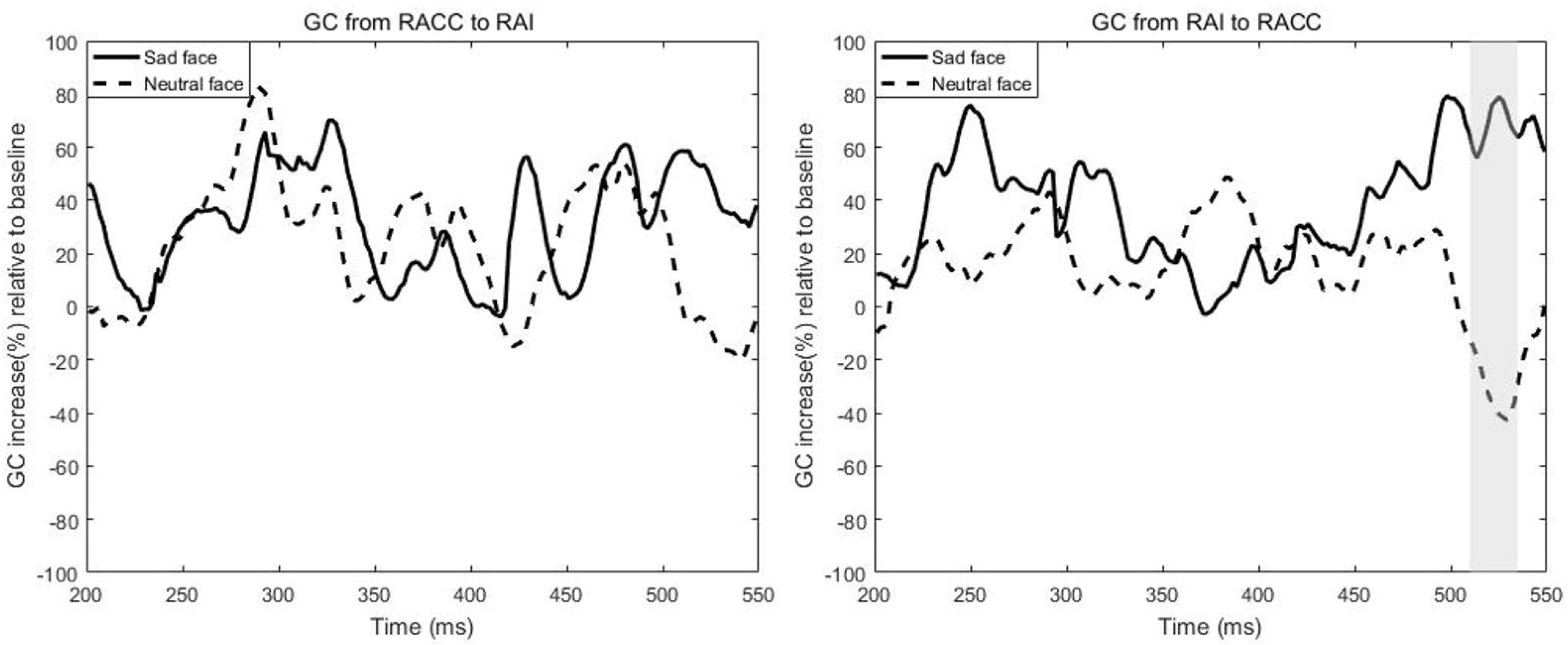
GC analysis results. There was increased bottom-up transfer of interoceptive information from the RAI to the RACC during sad face perception (left). The time in the figure represents the centre time of sliding time windows post-R peak, and the shaded area represents time windows that showed a significant difference between conditions after cluster corrections (Monte-Carlo p < 0.05, cluster-corrected). GCs from the [170 ms 230 ms] window to [425 ms 485 ms] window that were not included in the analysis are also plotted for visualization.

**Table 1.** Clusters showing significantly different time courses between sad face and neutral face conditions.

| Cluster | Region | MNI coordinates<br>(x, y, z) | Cluster size<br>(voxel number) | F value | Cluster P-value<br>(FWE-corrected) |
| --- | --- | --- | --- | --- | --- |
| Cluster 1 | Rt. SFG | 21, 20, 42 | 311 | 23.13 | 0.003 |
| **(Red)** | Rt. MFG | 33, 18, 36 |  | 12.86 |  |
| **Cluster 2** | Rt. AI | 27, 24, 6 | 379 | 20.02 | 0.001 |
| **(Green)** |  |  |  |  |  |
|  | Rt. putamen | 31, 6, 8 |  | 12.08 |  |
| **Cluster 3** | Rt. ACC | 9, 40, 10 | 313 | 17.32 | 0.003 |
| **(Blue)** |  |  |  |  |  |
|  | Lt. ACC | -3, 22, 28 |  | 14.68 |  |
| **Cluster 4** | Rt. putamen | 27, -8, 0 | 412 | 16.59 | <0.001 |
| **(Pink)** |  |  |  |  |  |
|  | Rt. GP | 21, 0, 2 |  | 15.30 |  |

SFG: superior frontal gyrus, AI: anterior insula, MFG: middle frontal gyrus, ACC: anterior cingulate cortex, GP: globus pallidus (voxel size: 1 mm x 1 mm x 1 mm)

#### GCs between sources of HEP modulations

The results of pairwise GC analysis showed that only the GC of the HEPs from the RAI to the RACC was significantly higher in the sad face condition than in the neutral face condition between 474 ms and 568 ms ([480 ms 540 ms] window to the [505 ms 565 ms] window, Monte-Carlo p = 0.014, cluster-corrected for 2 GCs and 27 time windows). The GC from the RACC to the RAI did not survive multiple comparisons (Monte-Carlo p = 0.146, cluster from [493 ms 553 ms] to [501 ms 561 ms]). These results indicate that only bottom-up information from the RAI to the RACC was increased.

### Analysis of physiological data

#### IBI and ECG R peak amplitude modulations in each condition

One-way repeated-measures ANOVAs showed no significant differences between conditions in either IBI (F (1.745, 54.095) = 1.513, p = 0.230, Greenhouse-Geisser-corrected, Fig. 5) or ECG R peak amplitude (F (2.989, 92.658) = 0.958, p = 0.416, Greenhouse-Geisser-corrected, Fig. 5).

**Fig. 5.**
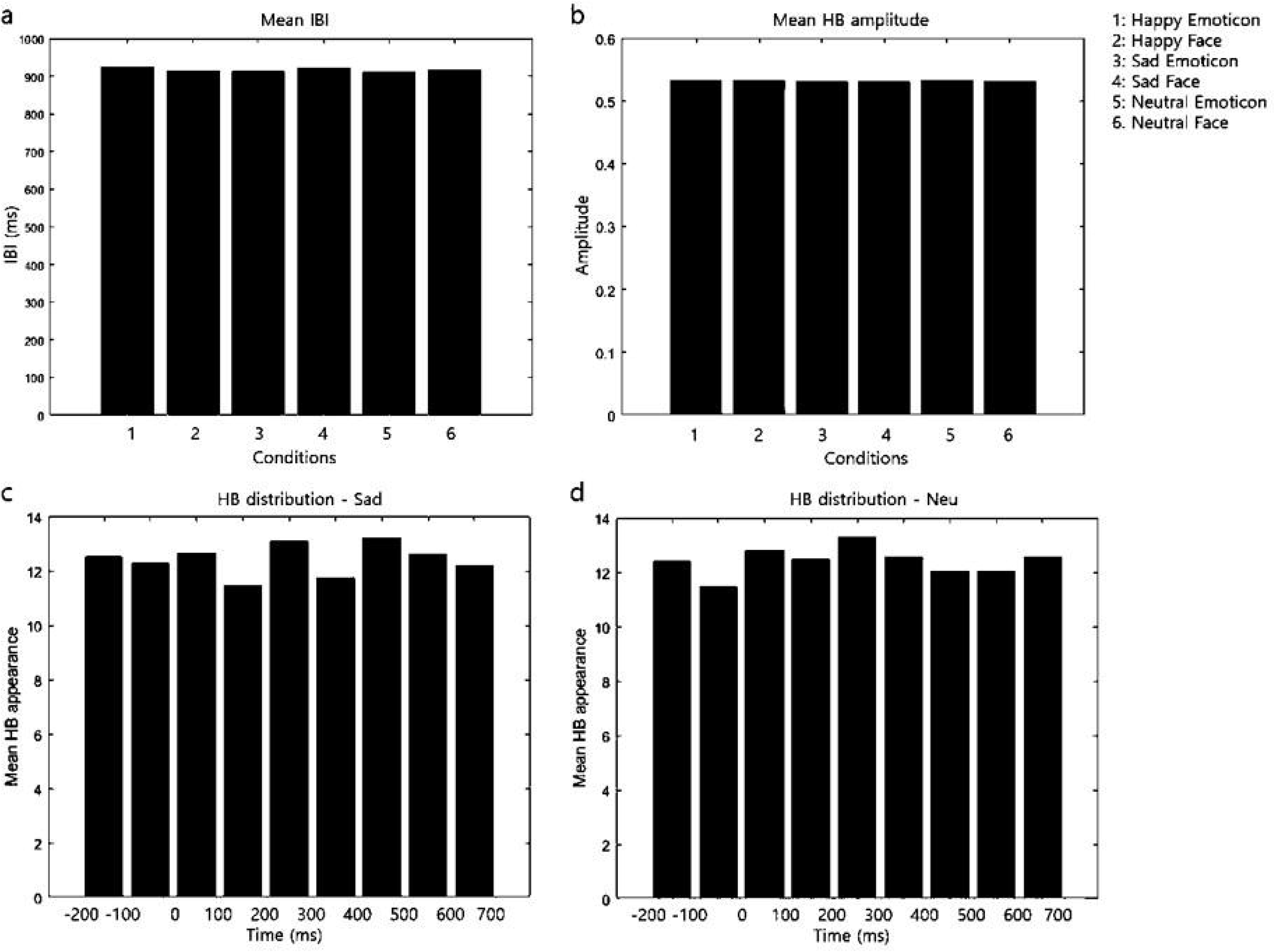
Physiological data analysis results. a. Mean IBIs of six experimental conditions. b. Mean R peak amplitudes of six experimental conditions, c. Mean R peak occurrence distributions in epochs of sad face (left) and neutral face (right) conditions.

#### Heartbeat distributions in each condition

The results of two-way 6*9 repeated-measures ANOVA (six conditions and nine 100-ms time bins (−200 ms to 700 ms from visual stimulus onset)) showed that there were no differences in the occurrence rate of heartbeats between the conditions (F (3.810, 118.111) = 0.783, p = 0. 533, Greenhouse-Geisser-corrected, Fig. 5) and no differences in the occurrence between time bins (F (4.876, 151.143) = 1.540, p = 0.182, Greenhouse-Geisser-corrected, Fig. 5).

### Analyses to exclude the effects of visual processing

#### Surrogate R peak analysis on the HEP modulation effect of a sad face

In the surrogate R peak analysis, our HEP modulation effect size (maximum cluster t statistics) was significant in the maximum cluster t statistics distribution of surrogate R peaks (Monte-Carlo p<0.03, Fig. 6), indicating that our effect was highly likely to be locked to the heartbeat.

**Fig. 6.**
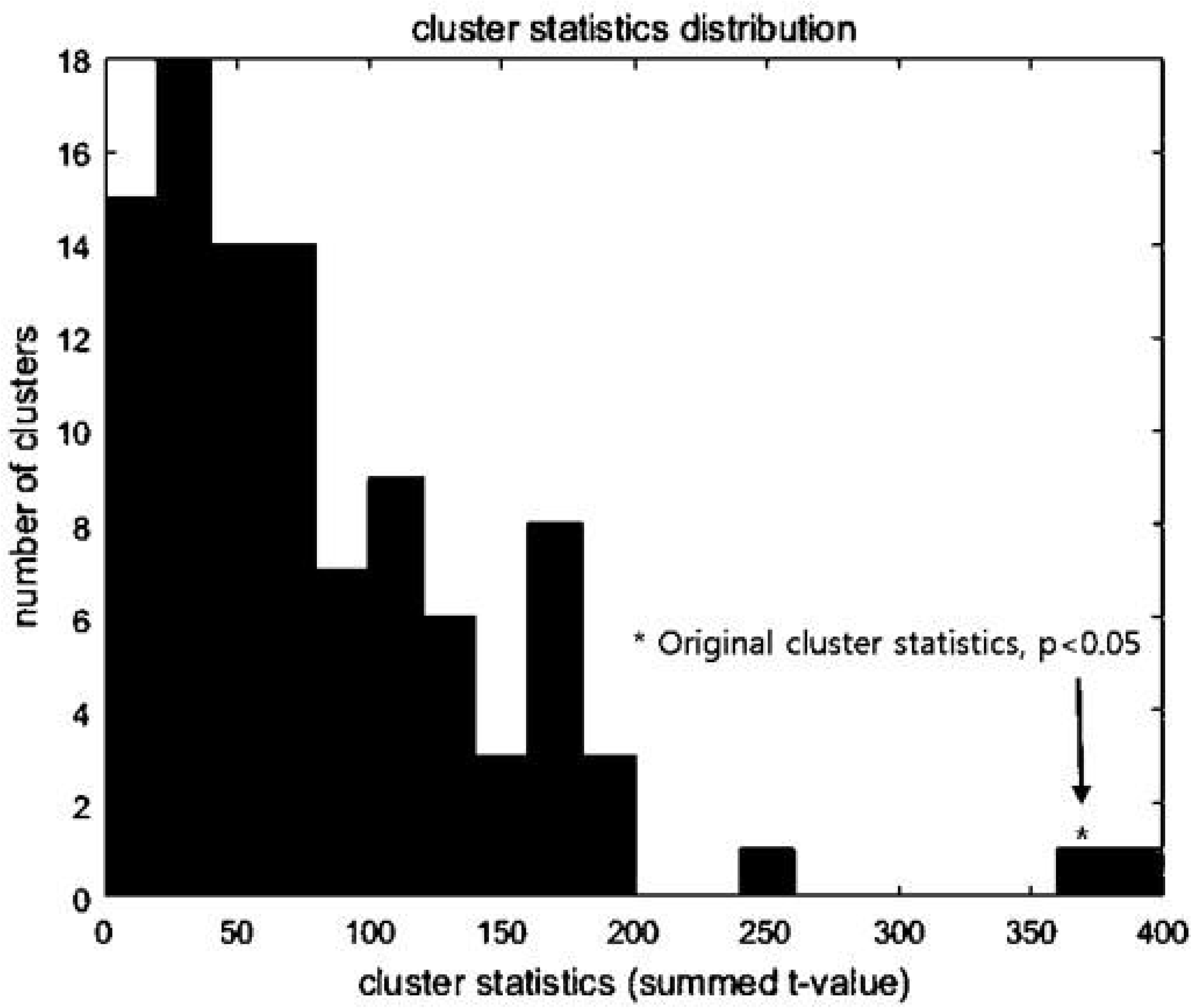
Surrogate R peak histogram (absolute cluster statistics). * represents the location of the original cluster statistics of the HEP modulation effect of a sad face.

#### Different spatial patterns between VEP and HEP analysis of sad face perceptions

In the cluster-based permutation *t* tests of VEPs comparing sad vs. neutral face conditions, two significant clusters were found at 73 ms-198 ms (Monte-Carlo p = 0.05) and 815 ms-948 ms (Monte-Carlo p = 0.004) after stimulus (Fig. S2). However, their topological distributions were completely different from those of the HEP effect. Furthermore, in the source analysis of VEPs (which was performed using the same method as the HEP source analysis), two clusters, which included a right ventromedial prefrontal cortex cluster and a right cuneus cluster, were found at 73 ms-198 ms (cluster-forming threshold = 0.01, minimum number of adjacent significant voxels = 10, cluster-level p-value < 0.001 in both clusters, Table 2, Fig. S3). Two clusters were also found in the 815 ms-948 ms time window, including a left cuneus cluster and a right cuneus cluster (p < 0.001 and p = 0.042, respectively, with the same cluster definitions of the earlier clusters, Table 2, Fig. S3). Figures describing the results of the sensor and source analyses of the VEPs are provided in the supplementary materials.

**Table 2.** Regions showing significantly different VEPs for a sad face compared to a neutral face (73 ms-198 ms, 815 ms-948 ms)

| **Cluster** | **Region** | **MNI** | **Cluster size** | **F value** | **Cluster P-** |
| --- | --- | --- | --- | --- | --- |
| **(73 ms-198 ms)** |  | **coordinates** | **(voxel** |  | **value (FWE-** |
|  |  | **(x, y, z)** | **number)** |  | **corrected)** |
| **Cluster 1 (Red)** | Rt superior | 11, 56, -17 | 576 | 18.18 | <0.001 |
|  | orbital gyrus |  |  |  |  |
|  | Rt medial orbital | 11, 36, -10 |  | 15.59 |  |
|  | gyrus |  |  |  |  |
|  | Lt gyrus rectus | -1, 50, -15 |  | 14.65 |  |

| **Cluster 2 (Blue)** | Rt cuneus | 3, -73, 18 | 465 | 18.17 | <0.001 |
| --- | --- | --- | --- | --- | --- |
| **Rt Cuneus** | Rt cuneus | 11, -68, 26 |  | 14.48 |  |
|  | Rt calcarine sulcus | 23, -48, 6 |  | 14.18 |  |
| **Cluster**<br>**(815 ms-948 ms)** | **Region** | **MNI**<br>**coordinates** | **Cluster size**<br>**(voxel**<br>**number)** | **F value** | **Cluster P-**<br>**value (FWE-**<br>**corrected)** |
| **Cluster 1 (Red)** | Lt calcarine sulcus | -3, -62, 14 | 836 | 33.02 | <0.001 |
|  | Lt cuneus | -1, -73, 20 |  | 25.06 |  |
|  | Lt calcarine sulcus | -1, -87, -4 |  | 23.78 |  |
| **Cluster 2 (Blue)** | Rt cuneus | 23, -68, 24 | 199 | 20.46 | 0.042 |
|  | Rt superior occipital gyrus | 28, -78, 34 |  | 15.89 |  |
|  | Rt superior occipital gyrus | 23, -73, 30 |  | 12.15 |  |

(voxel size: 1 mm x 1 mm x 1 mm)

## Discussion

Our findings provide direct evidence that the perception of a sad face modulates interoceptive information processing in the cortex.

First, we showed that cortical heartbeat processing after the presentation of a sad face has significantly different spatiotemporal dynamics than that of a neutral face and that these differences are localized to the interoceptive network (AI, ACC), basal ganglia (GP, putamen) and prefrontal areas (MFG, SFG). Importantly, the results of the GC analysis of these regions showed that bottom-up heartbeat information processing from the RAI to the RACC was increased in the sad face condition. In contrast to the HEP results, visual-locked activity was different in bilateral visual information processing areas including the bilateral cuneus and the ventromedial prefrontal cortex. Finally, surrogate R peak analysis provided strong evidence that our results were a consequence of cortical heartbeat processing modulation. Additionally, the analysis of physiological data and cardiac artefact removal using ICA also ruled out the possibility of other physiological effects on the cortical signal.

Our results extend previous studies of interoception and emotion in several aspects.

One pioneering study showed that emotional arousal modulates HEPs (Luft & Bhattacharya, 2015). Another recent study showed that HEPs are modulated even in infants by emotional stimuli including fear and angry facial expressions (Maister et al., 2017). Consistent with these studies, the sad face in our study also modulated HEPs, as seen with sensor analysis. However, the spatiotemporal patterns of HEP modulation in our sensor analysis were slightly different from these previous studies. The first study showed HEP modulation by emotional arousal in left parietal clusters at 305 to 360 ms after the R peak and in a right temporoparietal cluster at 380 to 460 ms after the R peak (Luft & Bhattacharya, 2015); in the second study, video clips of an angry or fearful facial expression increased HEPs at 150 to 300 ms after the R peak in a frontal cluster (Maister et al., 2017). However, considering the various spatiotemporal patterns found in HEP studies, these differences are not surprising. Additionally, the stimuli that were used in previous studies and in our study are different in some respects. Luft et al. showed an effect of emotional arousal by summing the arousal induced by both positive and negative emotional stimuli. In Maister et al., happy facial expressions did not show a HEP modulation effect, similar to the results of our study. However, angry and fearful expressions, which are negative emotions like sadness, showed spatiotemporal patterns (150-300 ms post-R peak in frontal clusters) of HEP modulation effects that are different from the current study. Because the subjects from this previous study were infants, whose IBIs are approximately half of those of adults, it is difficult to compare the temporal latencies between the results of that study and results of our study. However, the spatial pattern differences between the HEP modulation from Maister et al. and our study might be due to different emotion-induced interoceptive processing between anger/fear and sadness. Fear and anger are both high-arousal emotions, while sadness is a low-arousal emotion. Moreover, recent studies have reported that the perception of fearful expressions is influenced by the timing of the cardiac cycle, while the perception of a sad face stimulus is not affected (Garfinkel & Critchley, 2016); this finding shows that there are different interoception and emotion processing interaction patterns between fearful and sad faces. Considering these numerous differences, the differences in the spatiotemporal patterns of HEP modulation seen in the sensor analysis of our study and those of previous studies cannot be explained in a simple way. However, it is important to note that in addition to the previous studies showing HEP modulation using highly arousing emotional stimuli, our results also showed a HEP modulation effect in adults by the less arousing emotional stimuli of sad faces.

In the source analysis of HEP and VEP modulation by sad face perception, we found that there are two clearly distinct systems of processing that did not overlap, that is, cardiac information processing and visual information processing. For cardiac information processing, brain regions that reflect HEP modulations during sensor analysis were found in the RAI and the RACC, which are previously known sources of HEPs (Couto et al., 2015; H.-D. Park et al., 2017; H.-D. Park et al., 2014; Pollatos, Kirsch, & Schandry, 2005). These regions have also been identified as overlapping regions of emotion, interoception and social cognition processing in a recent meta-analysis (Adolfi et al., 2016). Moreover, a GC analysis revealed that cardiac information flow from the RAI to the RACC increases with sad face perception. The anterior insula interacts with many regions, including the ACC, which is a central autonomic network that regulates autonomic functions (Beissner, Meissner, Bär, & Napadow, 2013). More generally, these two regions are known to be main components of the salience network (SN) (Uddin, 2015). The detection and processing of salient stimuli are influenced by ascending interoceptive and visceromotor signals that indicate the moment-by-moment condition of the body and converge in the dorsal posterior insula (dPIC) and the mid-insula (Uddin, 2015). The SN, including the anterior insula and the anterior cingulate, integrates these ascending signals to coordinate large scale networks in the cortex (Uddin, 2015). In particular, the SN is known to play role in switching between the default mode network (DMN) and the central executive network (CEN). Moreover, the RAI to the RACC is proven to have a causal relationship in GC analysis using fMRI, while the reverse directional connectivity is weak (Sridharan, Levitin, & Menon, 2008). Considering these studies, our results of increased GCs of HEPs between the RAI and RACC might reflect increased emotion-related saliency processing induced by sad faces. However, whether this processing is specific to sad emotions or relevant to subjective feelings cannot be determined by our results. Further GC studies of HEPs using other emotional stimuli would clarify this point. In addition, recent studies have shown that neural responses to heartbeats in the DMN encode the two different dimensions of self-agentive ‘I’ and introspective ‘Me’ (Babo-Rebelo, Richter, et al., 2016; Babo-Rebelo, Wolpert, Adam, Hasboun, & Tallon-Baudry, 2016). In particular, heartbeat-evoked response (HER) difference between “high” and “low” ratings on the “I” scale were found in the ventral precuneus (vPC) in the 298-327-ms time window after the T wave (Babo-Rebelo, Richter, et al., 2016), and the amplitude of the HER was correlated with “I” ratings in the 444-500 ms after the R wave (Babo-Rebelo, Wolpert, et al., 2016). In our study, the HEP modulation effects were similar in the time windows of these studies. However, while increased information flow between SN regions and HEP modulation of the lateral prefrontal area including the SFG/MFG were identified, there was no HEP modulation within DMN-related regions in our results. From these observations, we suggest that the sad face-induced interoceptive information processing in the SN might reflect an orienting process to salient external emotional information rather than self-related information processing.

In the source analysis of VEPs, only visual cortical regions and ventromedial prefrontal regions were seen, which is consistent with previous EEG/MEG studies of sad faces (Batty & Taylor, 2003; Esslen, Pascual-Marqui, Hell, Kochi, & Lehmann, 2004). These regions did not overlap with the regions of HEP modulation. To our knowledge, this study is the first to show the distinctive processing of both systems induced by sad faces. These results correspond well to hypotheses that explain the relationship between emotions and interoception, such as the somatic marker hypothesis (A. R. Damasio, 2003), which predicts that bottom-up interoceptive processing is increased by emotional stimuli. However, note that the somatic marker hypothesis predicts that physiological changes are induced by emotional stimuli and modulate brain-body interactions, while the results of our physiological data analysis showed that cardiac activity parameters, including heart rate and heartbeat amplitude, were not modulated by sad face perception. However, considering that the direct input of the HEP is pressure in the carotid baroreceptor and that we did not measure blood pressure or indexes of carotid stimulation, it is difficult to determine whether there were physiological changes. Another possibility is that the absence of induced physiological changes might be due to our short SOA of approximately 1 second, during which it is difficult to evaluate physiological changes, such as heart rate modulation, that occur after several seconds (Critchley et al., 2005).

An unexpected finding in the present study was that cardiac information processing was also modulated in the basal ganglia (RGP/RP) and prefrontal regions (SFG/MFG). The GP is known to send input to the prefrontal cortex via the thalamus. This pathway is involved in initiating motor action (Singh-Bains, Waldvogel, & Faull, 2016). In particular, the ventral pallidum (VP) is closely involved in regulating emotion and initiating motor actions in response to emotional stimuli (Singh-Bains et al., 2016). Moreover, a case has been reported of a patient with damage to the GP, including the VP, who reported an inability to feel emotions (Vijayaraghavan, Vaidya, Humphreys, Beglinger, & Paradiso, 2008). Based on this evidence, we suggest that cardiac information is relayed to the GP (including the VP), and finally, the prefrontal area, which plays a role in initiating emotion-related behaviours such as facial expressions or the generation of feelings triggered by cardiac information processing; this processing is consistent with the somatic marker hypothesis.

Finally, note that we found HEP modulation to sad faces but not sad emoticons, which are also a sad emotional expression. We suspected that personal emotional experiences might influence interoceptive responses to sad emoticons. We provide and discuss some evidence towards this possibility following the additional analysis of sad emoticons (see supplementary materials).

To our knowledge, this study is the first to show different interoceptive and visual processing of the same emotional stimulus. In conclusion, our results demonstrate that the processing of sad faces induces different interoceptive information processing and visual processing compared to the processing of neutral faces, as reflected by HEPs and VEPs, respectively. Interoceptive processing involves increased bottom-up processing of the HEP from the RAI to the RACC, while visual processing occurs in different areas, including the visual cortical area. Additionally, we found that cardiac signals are also processed differently in the basal ganglia and prefrontal regions during sad face processing, which might reflect the initiation of emotion-related behaviours such as facial expression or the generation of feelings.

## Acknowledgements

This research was supported by the Brain Research Program through the National Research Foundation of Korea (NRF) funded by the Ministry of Science & ICT (NRF - 2016M3C7A1914448 and NRF - 2017M3C7A1031331 to B.J.) and partly supported by the BK21 plus fund (22A20151313464 to B.J.). The authors wish to acknowledge Kyung-Min An and Yong-Ho Lee for assistance with data acquisition. This manuscript was edited for English language by American Journal Experts (AJE).

