## Supplementary Materials for "Heartfelt Face Perception via the Interoceptive Pathway – an MEG study"

**Table S1. Inclusion criteria for GC ROIs**

| **ROI name** | **MNI coordinates (x, y, z)** |
| --- | --- |
| **RAI** | 35, 18, 4 |
| **RACC** | 9, 40, 10 |

ROIs were selected from peak voxels of a cluster, and a cluster was defined by an uncorrected p-value < 0.01 and the adjacent 10 voxels in the sad face – neutral face difference. RAI included 150 voxels, and RACC included 25 voxels (because RAI included more than 2 peak voxels).

**Additional analysis of the HEP modulation effect from a sad emoticon at the sensor level**

We attempted to determine why a sad emoticon, which is a sad emotional expression similar to a sad face, did not modulate HEPs although the emotionality scores for sad faces and sad emoticons were not different (paired t test, p = 0.492, mean emotionality score of sad emoticon = 2.81, mean emotionality score of sad face = 2.71). Unlike a face, a text-based emoticon is not an innately encoded stimulus, and responses to sad emoticons are likely to be acquired through experiences. Responses to emoticons may be affected by various factors including personal experiences, leading to variable responses to emoticons across the population. In our emotionality assessments, participants showed more heterogeneous responses to sad emoticons than to sad faces. The mean variance in the emotionality score for sad emoticons across subjects was 1.18, which was much larger than the variance of scores for sad faces (t (31) = 2.153, paired t test). Similarly, we hypothesized that the HEP modulation from a sad emoticon could be more variable than the HEP modulation from a sad face and could be more likely to be affected by other factors such as an individual’s emotional experience or their current mood. To prove this hypothesis, first, we attempted to determine the psychological traits or states that could modulate the HEP effect from a sad emoticon. We performed linear regressions with HEP modulation from a sad emoticon as the dependent variable and psychological variables as independent variables. Cluster-based permutation regression was performed using the FieldTrip function ‘ft_statfun_indepsamplesregrT’. We hypothesized that the HEP effect would occur in a time window similar to that of a sad face, while the spatial patterns could be different, based on our previous study of emoticons. In that study, the regions activated by emotional emoticons were not completely the same as the typical regions activated by emotional faces.

Therefore, the HEP values of time windows in which the HEP modulation effect was found for a sad face (488 ms-515 ms post-R peak) were averaged, while all sensors were tested and not averaged. Psychological variables used as independent variables included the PHQ-9 score, which reflects participants’ mood, and the VAQ score, which corresponds to a (negative) emotional experience. The VAQ included two subscales composed of peer verbal abuse history and parental verbal abuse history. Note that because two participants failed to complete the VAQ, only the data from thirty people were analysed.

Only the PeVA score showed a significant relationship with the HEP differences in the left central sensors (Monte-Carlo p = 0.015). Other psychological variables did not form significant clusters.

Based on this finding that HEP modulation is affected by PeVA history, we classified participants into two groups; the participants in the first group (n = 10) were those who had no peer verbal abuse at all, which correspond to a PeVA score of 15, while the second group consisted of those who had at least once incidence of peer verbal abuse (n = 20). Then, we performed a paired t test, which compared the averaged HEP for sad emoticons and neutral emoticons within the significant clusters found in the regression analysis for each group. A paired t test was also performed using the Monte-Carlo method because the number of participants in the first group was quite small. The results of the paired t test showed that there was a significant difference between the HEP from a sad emoticon and a neutral emoticon in the first group, which had no history of verbal abuse (Monte-Carlo p = 0.004, Fig. S4), while there was no difference in the second group, which had a verbal abuse history (Monte-Carlo p = 0.212). These results, showing both a significant relationship in HEP differences between sad and neutral emoticon conditions and the PeVA scores across all subjects and a significant difference in HEP differences in participants without a PeVA history suggest that the perception of a sad emoticon also modulates the interoceptive information processing that is influenced by subjective emotional experiences such as PeVA. Although these finding might not be a complete explanation for the lack of a HEP modulation effect from sad emoticons in contrast to the effect from sad faces, considering that there are very few studies related to the effects of emotional experiences on responses to emoticons, our results might provide clues to the effects of a person’s emotional experiences on their emotional response to acquired (not innate) emotional stimuli. Further studies of HEPs using various acquired emotional stimuli other than text-based emoticons such as Emojis are needed.

**Fig. S1. Examples of text-based emoticons.**

**Fig. S2. Topographic map (left figures) of the differences in VEPs between sad and neutral face conditions. Clusters showing significant difference are marked by black dots (Monte-Carlo p < 0.05, cluster-corrected). The clusters for an early time range (73 ms ~ 198 ms post stimulus) and a late time range are (815 ms ~ 948 ms) are plotted. Representative channel time courses of each cluster are plotted to the right of the topographic maps. The shaded areas represent the time points of clusters that showed a significantly different time course between conditions. Channels plotted in the right figures are marked with a white star in the topographic maps.**

**Fig. S3. Regions showing significantly different VEPs for a sad face compared to a neutral face (p < 0.01, cluster-level FWE-corrected, upper: 73 ms-198 ms, lower: 815 ms-948 ms).**

**Fig. S4. Topography of correlations between PeVA scores and HEP modulation by sad emoticons.**
