## Supplementary figures and images for "Heartfelt Face Perception via the Interoceptive Pathway – an MEG study"

### Supplementary Materials

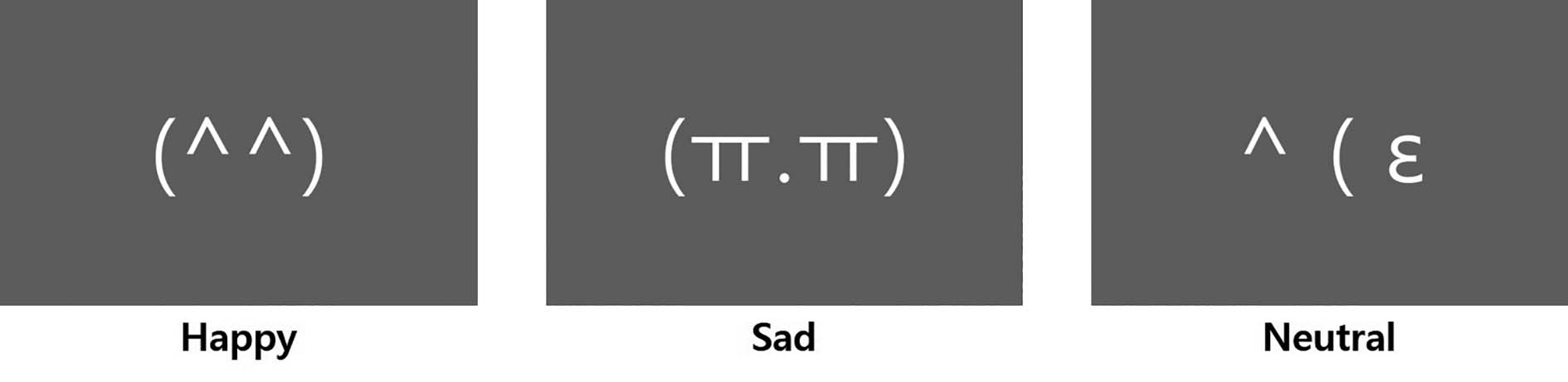

### Supplementary Materials

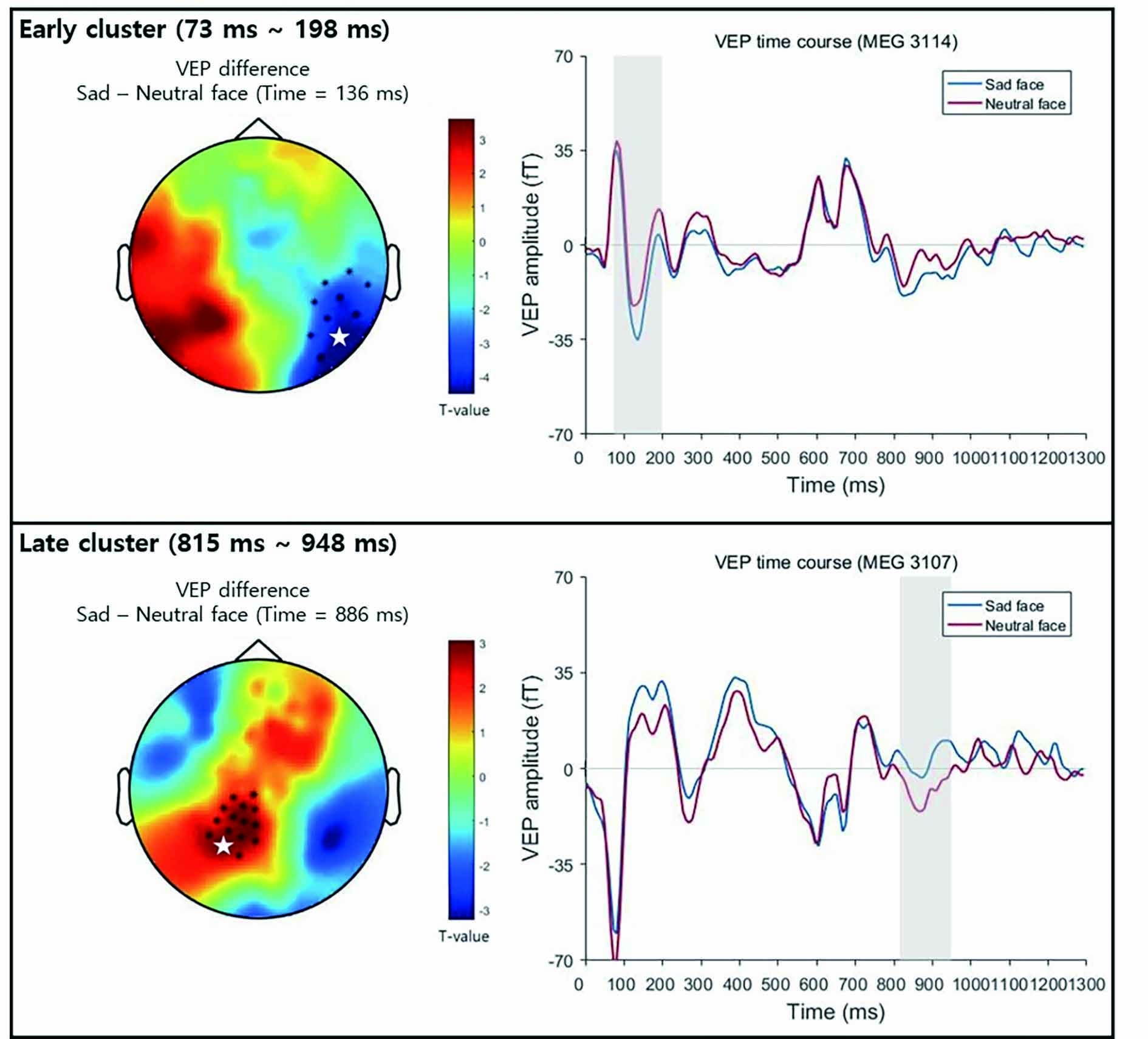

### Supplementary Materials

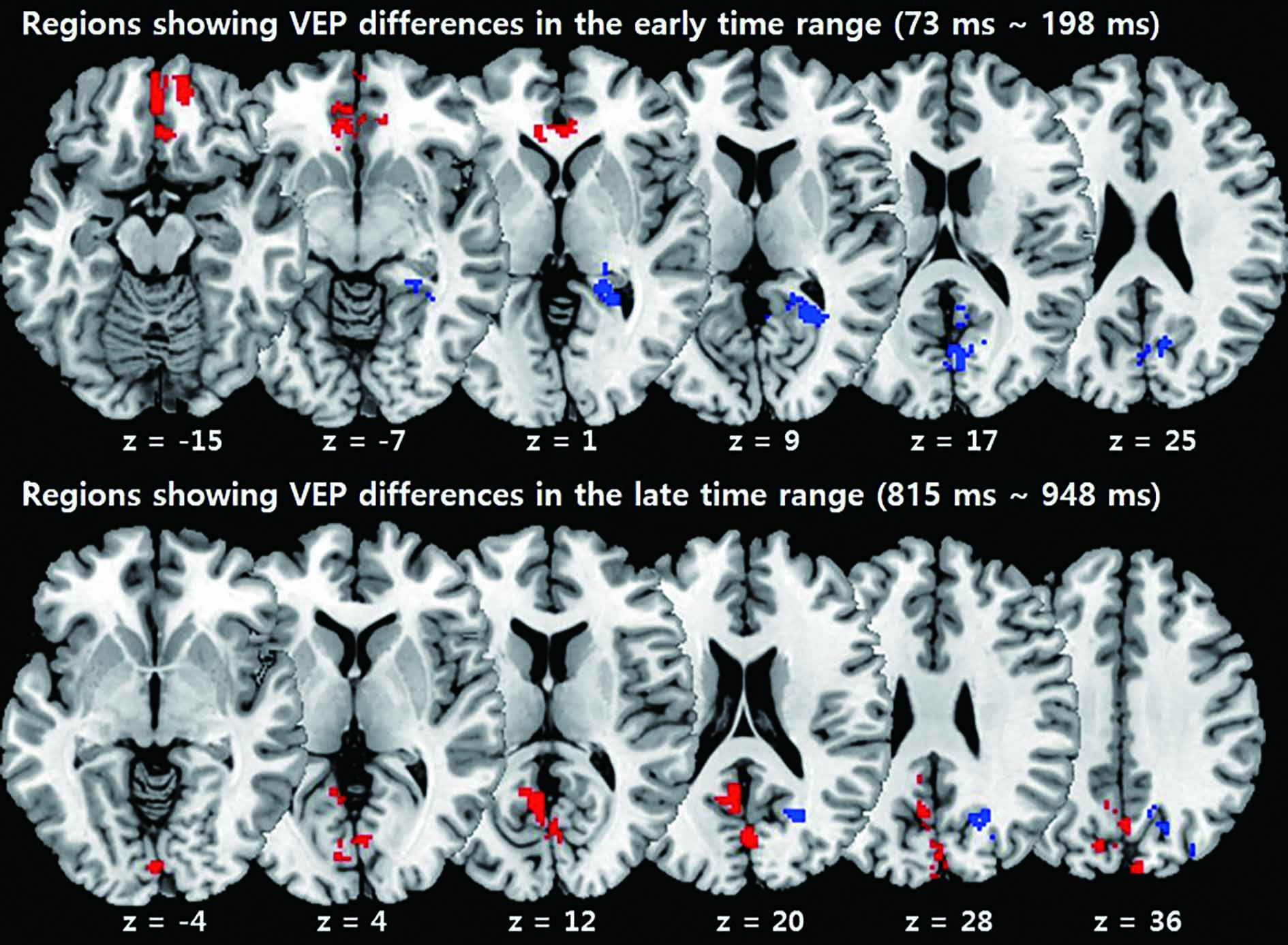

### Supplementary Materials

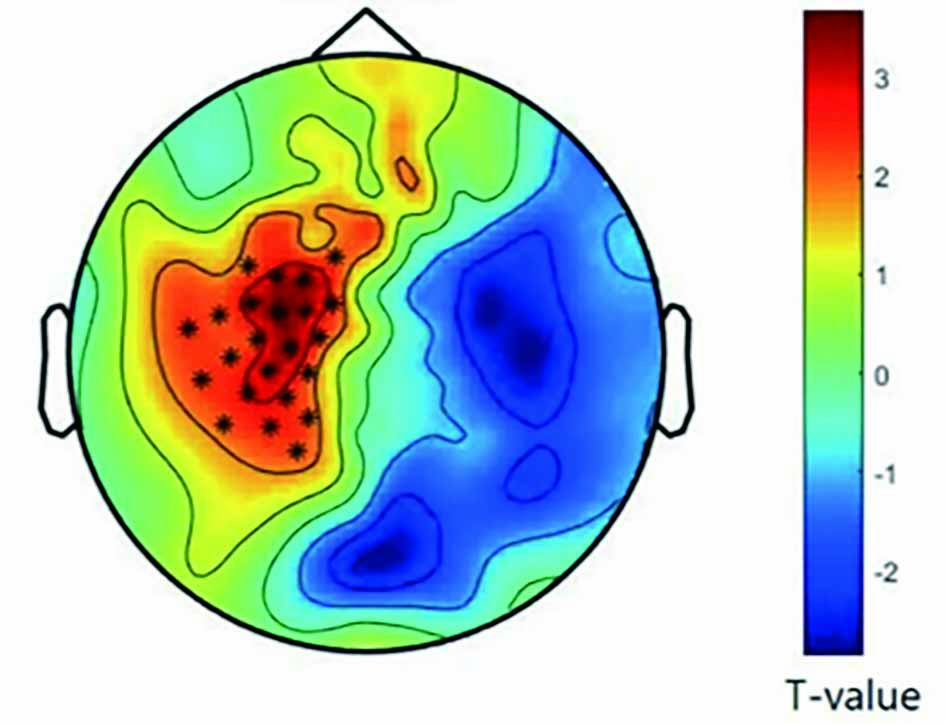
